# Induced pluripotent stem cell derived pericytes respond to endogenous mediators of proliferation and contractility

**DOI:** 10.1101/2023.09.29.560066

**Authors:** Natalie E. King, Jo-Maree Courtney, Lachlan S. Brown, Alastair J. Fortune, Nicholas B. Blackburn, Jessica L. Fletcher, Jake M. Cashion, Jana Talbot, Alice Pébay, Alex W. Hewitt, Gary P. Morris, Kaylene M. Young, Anthony L. Cook, Brad A. Sutherland

**Author notes:** Corresponding author: A/Prof. Brad A. Sutherland, Level 4, Medical Sciences Precinct, Tasmanian School of Medicine, University of Tasmania, Hobart, TAS 7000, Australia.

## Abstract

**Background:** Pericytes are multifunctional contractile cells that reside on capillaries. Pericytes are critical regulators of cerebral blood flow and blood-brain barrier function, and pericyte dysfunction may contribute to the pathophysiology of human neurological diseases including Alzheimers disease, multiple sclerosis, and stroke. Induced pluripotent stem cell (iPSC)-derived pericytes (iPericytes) are a promising tool for vascular research. However, it is unclear how iPericytes functionally compare to primary human brain vascular pericytes (HBVPs). We differentiated iPSCs into iPericytes of either the mesoderm or neural crest lineage using established protocols. We compared iPericyte and HBVP morphologies, quantified gene expression by qPCR and bulk RNA sequencing, and visualised pericyte protein markers by immunocytochemistry. To determine whether the gene expression of neural crest iPericytes, mesoderm iPericytes or HBVPs correlated with their functional characteristics *in vitro*, we quantified EdU incorporation following exposure to the key pericyte mitogen, platelet derived growth factor (PDGF)-BB and, contraction and relaxation in response to the vasoconstrictor endothelin-1 or vasodilator adenosine, respectively. iPericytes were morphologically similar to HBVPs and expressed canonical pericyte markers. However, iPericytes had 1864 differentially expressed genes compared to HBVPs, while there were 797 genes differentially expressed between neural crest and mesoderm iPericytes. Consistent with the ability of HBVPs to respond to PDGF-BB signalling, PDGF-BB enhanced and PDGF receptor-beta inhibitors impaired iPericyte proliferation. Administration of endothelin-1 led to iPericyte contraction and adenosine led to iPericyte relaxation, of a magnitude similar to the response evoked in HBVPs. We determined that neural crest iPericytes were less susceptible to PDGFR beta inhibition, but responded most robustly to vasoconstrictive meditators. iPericytes express pericyte-associated genes and proteins and, exhibit an appropriate physiological response upon exposure to a key endogenous mitogen or vasoactive mediators. Therefore, the generation of functional iPericytes would be suitable for use in future investigations exploring pericyte function or dysfunction in neurological diseases.

## Background

Pericytes are contractile cells that reside within the capillary bed. In the cerebrovasculature, pericytes are essential regulators of cerebral blood flow and contribute to blood-brain barrier formation and function (1). Pericyte dysfunction may contribute to the pathophysiology of neurological diseases including Alzheimer’s disease, stroke, and multiple sclerosis (1, 2). For example, the aggregation of amyloid-β, a key protein associated with Alzheimer’s disease pathology, induces pericyte constriction by modulating the endothelin-1 receptor signalling pathway (3). Furthermore, pericytes die during stroke, in a way that constricts capillaries and prevents tissue reperfusion even after large vessels reopen – a phenomenon known as ‘no-reflow’ (4). In addition, pericytes can also have reparative properties as it has been shown that activation of the pericyte PDGFRβ signalling pathway can facilitate repair following a stroke, by supporting fibrotic scar formation (5).

iPSC-derived pericytes (iPericytes) are increasingly used in place of primary pericyte lines to model pericyte function in health and disease (6–11). iPericytes have several advantages over primary pericyte lines, as they can be derived from iPSCs reprogrammed from individuals of various genetic backgrounds and disease diagnoses (12), allowing them to be used for basic biological studies as well as disease modelling or phenotyping. It is also possible to co-culture iPericytes with cells derived from the same iPSC line, to model the neurovascular unit (NVU) (13). Finally, iPericytes may be compatible with personalised medicine approaches as they could be returned to the donor without immune rejection – an approach that was recently validated in mice (14).

There are several published methods for iPericyte differentiation (7–11, 15). One describes a 10-day method for generating iPericytes of two developmental lineages: neural crest or mesoderm iPericytes (16). The iPericytes had morphological features that were consistent with primary pericyte lines and expressed key pericyte markers including PDGFRβ, alanyl aminopeptidase (CD13) and neuron-glial antigen 2 (NG2) (16). While the expression of key pericyte markers is promising, it is essential to understand the functional capacity of iPericytes relative to primary pericyte lines. iPericytes can increase endothelial cell expression of BBB markers in co-culture, improve trans-endothelial electrical resistance (TEER) and enhance the formation of 3D endothelial cell tubes (7, 9, 10, 15, 16). However, the proliferative and contractile functions of iPericytes have not been explored, and a side-by-side comparison of iPericytes and primary HBVPs is also lacking.

In this study, we therefore aimed to characterise the gene expression profiles, and proliferative and contractile properties of neural crest and mesoderm iPericytes and compared them to HBVPs. We compared the PDGF-BB and PDGFRβ-mediated mitogenic response of neural crest and mesoderm iPericytes, and quantified cell area change in response to the vasoconstrictor, endothelin-1 and vasodilator, adenosine. We report that iPericytes have functional PDGFRβ signalling, capable of mediating proliferation. Furthermore, iPericyte area changes in response to endothelin-1 and adenosine. iPericytes are functionally similar to HBVPs, making them suitable for use in *in vitro* assays and for disease modelling.

## Materials and Methods

### Pluripotent Stem Cell Lines

The TOB-00220 iPSC line (from a 67 year-old male apparently healthy donor) (17) was cultured to generate mesoderm and neural crest iPericytes with approval from the University of Tasmania Human Research Ethics Committee (Project H26563). Additional healthy control iPSC lines were used as specified in text: MNZTASi019-A (from a 53 year-old female donor); MNZTASi021-A (76 year-old male donor), and MNZTASi022-A (56 year-old female donor) were purchased from the MS Stem biobank (Menzies Institute for Medical Research, Hobart, Tasmania, Australia). MS Stem iPSCs were generated and characterised as previously described (18, 19) with approval from the University of Tasmania Human Research Ethics Committee (Project H16915). All iPSC lines were shown to have karyotypically normal karyograms within 10 passages of use for experiments and were used between passage 5-35.

### Pluripotent Stem Cell Culture

iPSCs were grown on Matrigel (Merck, cat.#354277) coated plates in mTeSR+ cell culture medium (Stem Cell Technologies, cat.#05825) maintained at 37°C in a 20% O_2_ / 5% CO_2_ humidified incubator. The culture medium was exchanged every 2 days, and iPSCs were cultured to generate large colonies (∼60-100µm diameter) with distinct round edges. iPSC colonies were passaged using Versene Solution (Gibco, cat.#15040066).

### Differentiation of iPSCs into mesoderm or neural crest iPericytes

iPSCs were differentiated to produce iPericytes by adapting a previously published protocol (16) (see Fig. S1). Induction into mesoderm iPericytes was achieved by culturing in Mesoderm Induction Media (Stem Cell Technologies, cat.#05221). Induction into neural crest iPericytes was achieved by culturing in DMEM/F-12 plus GlutaMAX (Thermofisher Scientific, cat.#10565018) supplemented with 0.5% (v/v) Bovine Serum Albumin (Sigma Aldrich, cat.#A9418), 2% (v/v) B27 (ThermoFisher Scientific, cat.#17504-044) and 3µM CHIR 99021 (GSK3 inhibitor; Tocris Bioscience, cat.#TB4423-GMP). Medium was exchanged daily for 5 days before it was replaced with Complete Pericyte Medium (CPM, ScienCell Research Laboratories, cat.#1201), which was exchanged daily for a further 5 days. After 10-days, the resulting iPericytes were maintained as outlined below.

### Pericyte culture

Human brain vascular pericytes (HBVPs, ScienCell, cat.#1200) and iPericytes were grown in CPM which was replaced every second day. Pericytes were passaged at 60%-90% confluence by washing with Dulbecco’s phosphate buffered saline without magnesium or calcium (DPBS^-/-^, ThermoFisher Scientific, cat.#14190-144) prior to treatment with TrypLE Express (Thermofisher Scientific, cat.# 12604013). Pericytes were passaged and the cells allowed to adhere for ≥ 16h prior to commencing experiments. All HBVPs and iPericytes were used between passage 2-8.

### Real Time qPCR

To quantify the expression of pericyte-associated genes in HBVPs, iPSCs, mesoderm iPericytes or neural crest iPericytes using real time quantitative polymerase chain reaction (qPCR), cells were grown to 95% confluence in 6-well plates (Interpath, cat.#657160). Cells were collected from n = 3 wells per cell type of the same differentiation, and RNA was extracted using an RNeasy mini kit (Qiagen,cat.#74104), following the manufacturer’s recommendations. RNA concentration was quantified using a NanoDrop (ND-1000, Thermofisher Scientific) and RNA quality was evaluated in a subset of samples using an Agilent 4200 Tape Station system (cat.#G2991AA) with an RNA ScreenTape Ladder (Agilent, cat.#5067-5578), following the manufacturer’s instructions. cDNA synthesis was performed using the High Capacity cDNA Reverse Transcription Kit (Thermofisher Scientific, cat.#4368814). For reverse transcription a SuperCycler Trinity (Kyratec, cat.#SC-200) was set to the program: step 1-25°C, 10 min; step 2 - 37°C, 120min; step 3 - 85°C, 5 min; step 4 – 4°C, infinity. 200ng of cDNA was added to the TaqMan Fast Advanced Master Mix (Thermofisher Scientific, cat.#4444557) and TaqMan primers for mRNAs of interest for each 20 µL qPCR reaction. MicroAmp Optical 96 Well Reaction Plates (Thermofisher, cat.#N8010560) were placed in a QuantStudio 3 (Thermofisher Scientific, cat.#A28567) operating the following program: step 1-50°C, 2 min; step 2 - 95°C, 2 min; step 3 - 95°C, 1 sec then 60°C, 20 sec (X 40). Raw data were exported into the QuantStudio Design and Analysis Software (v1.5.1, Applied Biosystems) to calculate Cycle threshold (Ct) values for each sample. Delta Ct values, delta delta Ct values and 2^-delta delta Ct were calculated in Microsoft excel, using *HPRT1* as a housekeeping gene. Primers included: *CSPG4* (Hs00361541_g1, Thermofisher Scientific, cat.# 4331182), *OCT4* (Hs01895061_u1, Thermofisher Scientific, cat.# 4331182), *NANOG* (Hs04399610_g1, Thermofisher Scientific, cat.# 4331182), *ACTA2* (Hs00426835_g1, Thermofisher Scientific, cat. #4331182), *PDGFRB* (Hs01019589_m1, Thermofisher Scientific, cat.# 4331182) and *HPRT1* (Hs02800695_m1, Thermofisher Scientific, cat.# 4331182).

### Immunocytochemistry

For immunocytochemical studies, HBVPs, mesoderm iPericytes and neural crest iPericytes were plated in Greiner 24 Well Plates (Interpath, cat.#662160X) and grown to 50% confluency. Medium was removed and cells were fixed by immersion in ice-cold methanol (100%) for 10 min prior to washing with ice-cold PBS (Gibco, cat.#18912014). Cells were washed thrice with 0.1% (v/v) tween-20 / PBS and incubated in Serum Free Protein Block (DAKO, cat.#X0909) for 1h at 21°C. Primary antibodies (rabbit anti-CD13, Abcam Ab108310, RRID:AB_10866195; rabbit anti-NG2, Sigma Aldrich AB5320, RRID:AB_91789; rabbit anti-PDGFRβ, Abcam Ab32570, RRID:AB_777165; rabbit anti-αSMA, Abcam Ab5694, RRID:AB_2223021) were diluted 1:200 in Antibody Diluent (DAKO, cat.#S302283-2) and applied to cells overnight at 4°C. Cells were washed thrice in 0.1% (v/v) tween-20 / PBS before applying secondary antibody (Alexa Fluor 488-conjugated Donkey anti-rabbit, ThermoFisher Scientific, cat.# A-21206) diluted 1:1000 in Antibody Diluent for 2 h at 21°C in the dark. Cells were washed thrice in PBS and incubated with 4′,6-diamidino-2-phenylindole (DAPI) (Sigma, #D9542) diluted 1:10,000 in PBS for 5 min. Cells were imaged at 10x using a Cytation 5 Cell Imaging Multi-Mode Reader (Biotek, USA).

### RNA sequencing, data processing and differential gene expression analysis

Samples containing > 10 ng/µl RNA with a RIN of > 8 were sent to the Australian Genome Research Facility for bulk RNA sequencing. Libraries were generated using an Illumina Stranded mRNA workflow with polyA capture. RNA sequencing, processing of raw sequencing data, and quantification of gene expression are described in the supplementary methods. Differential gene analysis, principal components analysis (PCA), gene ontology analysis and heatmap generation were performed using DESeq2 and other tools as described in supplementary methods.

### Proliferation Assay

An EdU assay was used, as described previously (20), to quantify proliferation in mesoderm or neural crest iPericytes compared to HBVPs. Briefly, pericytes were plated in 96 well plates (Interpath, cat.#655180) and grown to 50% confluency (∼5,000 cells per well). CPM was replaced with either: CPM containing the complete array of pericyte growth factors, incomplete pericyte media (PM) which did not contain any growth factors, PM supplemented with 100 ng/mL PDGF-BB (Sigma Aldrich, SRP3138) or PM supplemented with 100 ng/mL PDGF-BB with either 0.1 µM, 10 µM or 100 µM imatinib (Sapphire Bioscience, 00022120). Pericytes were cultured for 24 h prior to fixation by immersion in 4% (w/v) PFA in PBS for 15 mins at 21°C. EdU incorporation into the DNA was revealed using a Click-iT EdU Cell Proliferation Kit (Invitrogen, cat.#C10340) following the manufacturer’s instructions, and the nuclei of all cells were identified by staining with DAPI. EdU and DAPI labelling was visualised and imaged at 20x magnification using a Nikon Ti2 SRRF microscope. A region of interest spanning 3 mm^2^ (20x magnification, 3×3) was defined, imaged and stitched to create a single image spanning the region of interest for quantification. QuPath V0.2.3 was used to identify total cells from the DAPI channel as well as proliferative cells from the EdU channel using techniques previously described (21). Briefly, channel colours (DAPI, EdU) were set for all images as a batch using the script “Channels and colours.groovy”, described previously (21). The rectangle annotation tool was used to draw a ROI around each image using the script “Select all ROI.groovy”. DAPI and EdU positive cells were detected using the Positive Cell Detection tool using the script “EDU Analysis”. Proliferation was calculated as:

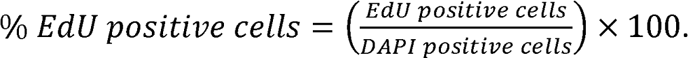

### Contraction Assay

An xCelligence Real-Time cell analysis electrical impedance assay was used, as previously published (22), to quantify contractility in mesoderm or neural crest iPericytes compared to HBVPs. 5,000 pericytes were plated in each well of an E-Plate (ACEA Biosciences, cat.# 05469830001), with 200 µL of CPM. After ∼16 h, cells were above 50% confluence and CPM was replaced with CPM alone (control) or CPM containing 50 nM endothelin-1 or 10 µM adenosine, concentrations as used previously (22, 23) (n = 4 wells per condition) and the plate was placed in the xCelligence system. The xCelligence system measures the relative impedance of electron flow expressed as arbitrary ‘cell index’ units as an indicator of cell area (Fig. S4). Cell index was measured every minute for 2 h at 37°C and 5% CO_2_. The normalised cell index value was calculated by normalising the raw cell index values to the cell index value at baseline t=0 as described previously (24). Area under the curve (AUC) was calculated using GraphPad prism for the normalised cell index graphed over the first 20 mins following drug exposure. Change (Δ) in cell index was calculated at the maximum point of contraction in each well (Maximum Δ Cell Index) using the equation:

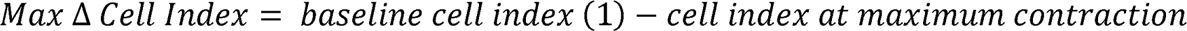

Change in cell index was also calculated after 2h (Δ Cell index after 2 h) to determine the maintenance of contraction after 2 h using the

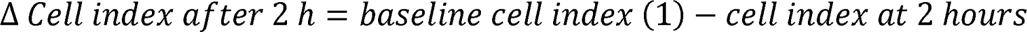

See Fig. S4 for more details about the xCelligence system and calculations.

### Statistical Analyses

Statistical analyses were performed using Prism 9.3.1 (GraphPad, USA) except for RNA-seq data where DESeq2 and R were used (see supplementary methods for details about RNA-seq analysis). Prior to performing statistical comparisons in Prism, outliers were removed using the ROUT’s outlier test (*Q* = 1%). Each data set was tested for normality of residuals using the Shapiro-Wilk test, and either a Y=Log(Y) transformation was performed to enable parametric testing, or data sets were analysed with non-parametric Mann-Whitney U or Kruskal-Wallis tests. To compare qPCR data generated from iPSCs and iPericytes, we performed a one-way ANOVA with a Dunnett’s multiple comparisons test or Sidak’s multiple comparisons test. To determine the effect of experimental conditions on proliferation, we performed a one-way ANOVA, with differences between conditions versus control determined using a Dunnett’s multiple comparisons test. For the contraction assay, we performed a two-way ANOVA to determine the effect of cell type (mesoderm iPericytes, neural crest iPericytes, or HBVPs) or treatment (control, endothelin-1 or adenosine) on cell index parameters, followed by a Tukey’s multiple comparison test for pair-wise comparisons. A *p* < .05 was considered statistically significant. Statistical tests and results for each analysis are reported in the figure legends.

## Results

### iPericytes display characteristic pericyte morphology and express canonical pericyte markers

To determine whether iPericytes have the morphological characteristics of pericytes, we collected phase contrast micrographs of mesoderm and neural crest iPericytes and HBVPs and assessed the morphological features of each cell type. Mesoderm and neural crest iPericytes had elongated fusiform cell bodies, that were similar in morphology to HBVPs (Fig. 1A). *In vitro,* HBVPs adopt several morphological phenotypes, that relate to different contractile “subsets” (23). Mesoderm and neural crest iPericytes cultures also contained each of these morphological phenotypes (Fig. S2) and in proportions similar to those reported for HBVPs (23). To determine whether iPericytes express classical pericyte markers, we isolated RNA and generated cDNA to conduct a qPCR analysis. iPericytes expressed mRNAs that are integral to pericyte function, particularly: *PDGFRB*, which encodes the PDGFRβ protein; *CSPG4* which encodes NG2 proteoglycan, and *ACTA2* which encodes alpha-smooth muscle actin (αSMA). Compared to iPSCs, HBVPs, mesoderm and neural crest iPericytes expressed significantly higher levels of *CSPG4* (HBVP, p = 0.0009; neural crest iPericytes, p < 0.0001; mesoderm iPericytes, p < 0.0001), *ACTA2* (HBVP, p = 0.0489; neural crest iPericytes, p = 0.0022; mesoderm iPericytes, p = 0.0190), and *PDGFRB* (HBVP, p = 0.0947; neural crest iPericytes, p < 0.0001, mesoderm iPericytes, p = 0.0002) mRNA (Fig. 1B). Conversely, HBVPs, neural crest and mesoderm iPericytes expressed pluripotency genes at a very low level; expressing less *OCT4* (HBVP, p < 0.0001; neural crest iPericyte, p < 0.0001; mesoderm iPericyte, p < 0.0001) and *NANOG* (HBVP, p < 0.0001; neural crest iPericyte, p < 0.0001; mesoderm iPericyte, p < 0.0001) mRNA than iPSCs (Fig.1B).

**Figure 1.**
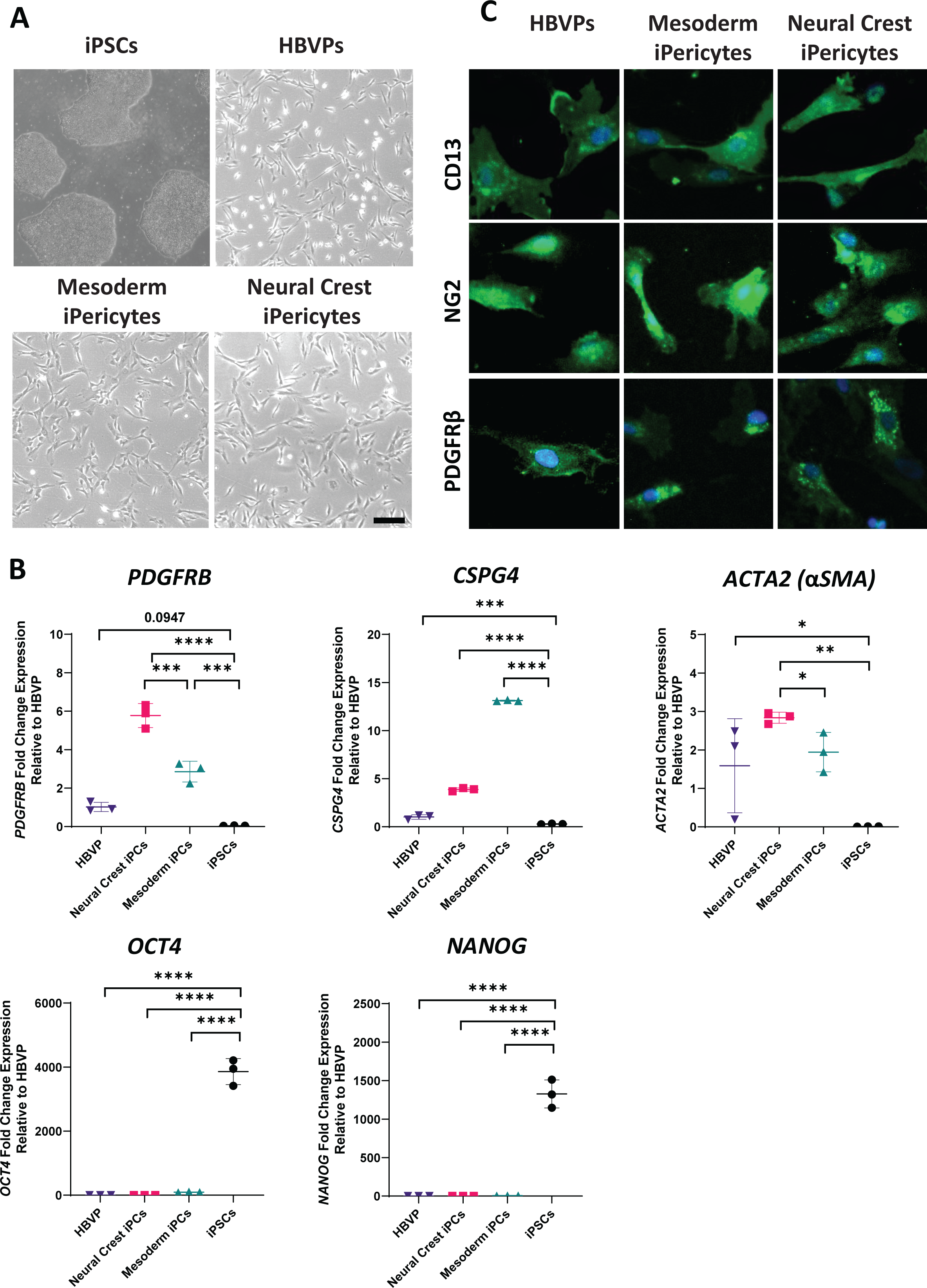
iPericytes are morphologically similar to HBVPs and express pericyte markers. (A) Phase contrast bright 4x magnification images of iPSCs, HBVPs, mesoderm iPericytes and neural crest iPericytes. Scale = 200µm. (B) Fold change gene expression measured by qPCR of pericyte genes *PDGFRB, CSPG4, ACTA2* and pluripotency genes *OCT4 and NANOG* by iPSCs, neural crest iPericytes, mesoderm iPericytes and HBVPs (n = 3 per cell type). Data are normalised to HBVP cells, and comparisons were made using a one-way ANOVA: *PDGFRB* (F (3, 8) = 103.1, p < 0.0001), *CSPG4* (F (3, 8) = 4671, p < 0.0001), *ACTA2* (F (3, 8) = 9.340, p < 0.0054), *OCT4* (F (3, 8) = 1686, p < 0.0001) and *NANOG* (F (3, 8) = 606.4, p < 0.0001). Post-hoc comparisons performed using Dunnett’s multiple comparisons test: * p < 0.05, ** p < 0.01, *** p < 0.001,**** p < 0.0001. Data are shown as mean ± SD. (C) Immunocytochemistry showing expression of pericyte proteins CD13, NG2 and PDGFRβ (green) by HBVP, mesoderm iPericytes and neural crest iPericytes. Nuclei counter-stained with DAPI (blue). Scale = 20 µm.

To extend these mRNA expression findings, we performed immunocytochemistry to determine whether iPericytes expressed proteins synonymous with pericyte identity: CD13, NG2 and PDGFRβ. Mesoderm and neural crest iPericytes displayed a pattern of labelling which indicated that proteins were expressed within similar sub-cellular locations with anti-CD13, anti-NG2 and anti-PDGFRβ antibodies compared to HBVPs (Fig. 1C). Overall, these data show that iPericytes are morphologically similar to HBVPs and express mRNAs and proteins that are consistent with pericyte identity.

### Mesoderm and neural crest iPericytes have different gene expression profiles

To identify differences in gene expression between mesoderm and neural crest iPericytes, and to determine how similar these cells are to HBVPs, we performed bulk RNA sequencing. A PCA revealed that the majority of the variance was accounted for through the difference between HBVPs and iPericytes regardless of lineage (PC1: 78% variance), whereas PC2 (12% variance) accounted for the variation between neural crest and mesoderm iPericytes (Fig. 2A). Differential gene expression analysis was used to explore differences between HBVPs and iPericytes (Fig. 2B-D), or neural crest and mesoderm iPericytes (Fig. 2E-G). There were a substantial number of differentially expressed genes between HBVPs and iPericytes, with 984 genes upregulated and 880 genes downregulated in iPericytes compared to HBVP (Fig. 2B). This is also reflected in the heat map with clear differences in gene expression between HBVPs and iPericytes, regardless of lineage (Fig. 2C). Gene ontology analysis of differentially expressed genes between HBVPs and iPericytes showed enrichment for genes related to tissue development, cellular division, morphology, extracellular matrix production and protein binding (Fig. 2D).

**Figure 2.**
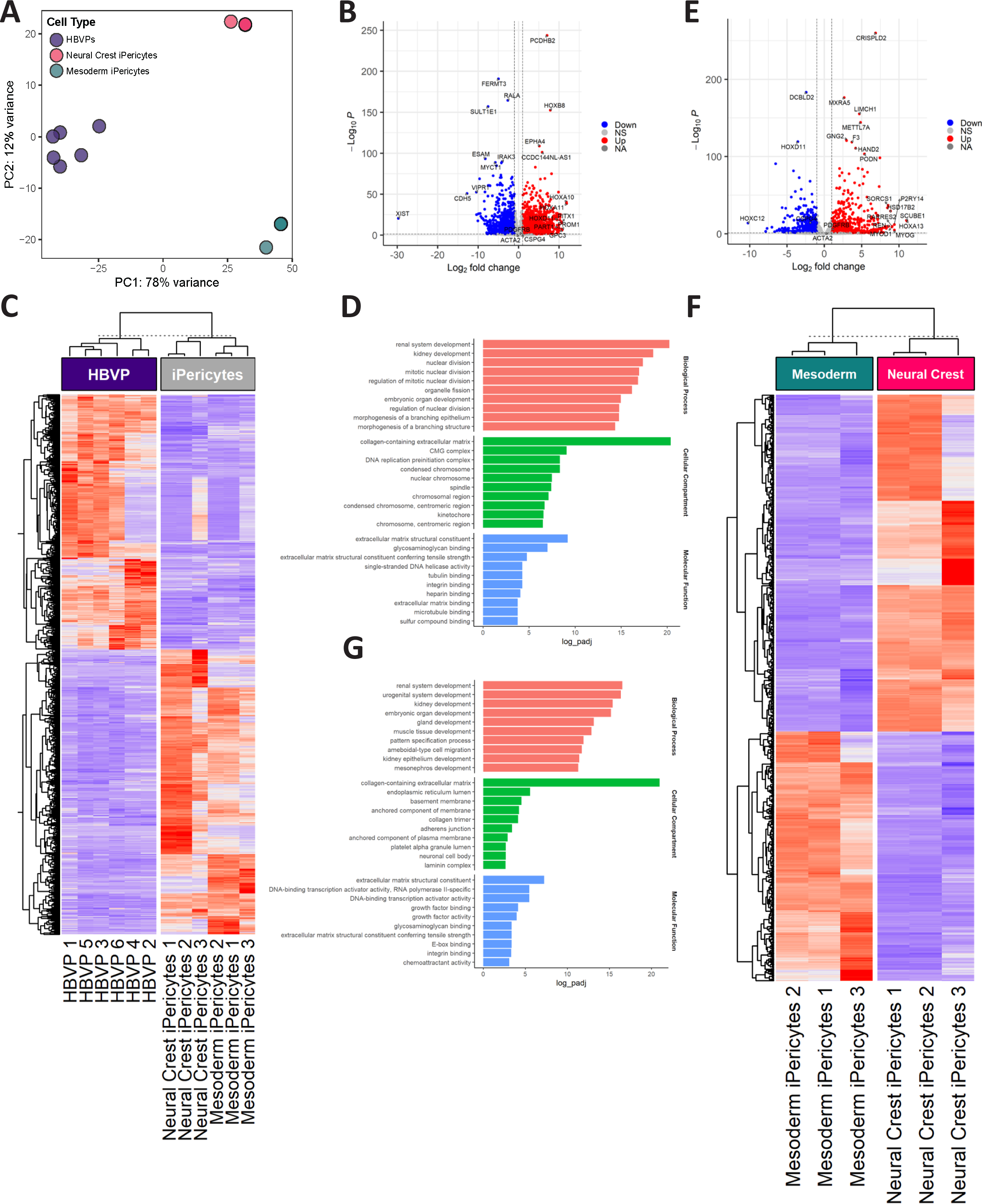
iPericytes derived through different lineage pathways have differential expression of genes. (A) PCA analysis showing separate clustering of mesoderm iPericytes, neural crest iPericytes and HBVPs (n = 6 for HBVPs, n = 3 for mesoderm or neural crest iPericytes). (B) Volcano plots showing upregulated and downregulated genes in iPericytes compared to HBVPs. (C) Heat map showing differentially expressed genes in iPericytes compared to HBVPs. (D) Gene ontology analysis of key biological processes, cellular compartments and molecular function associated with 1,864 differentially expressed genes between iPericytes and HBVPs. (E) Volcano plots showing upregulated and downregulated genes in neural crest iPericytes compared to mesoderm iPericytes. (F) Heat map showing differentially expressed genes in neural crest iPericytes compared to mesoderm iPericytes. (G) Gene ontology analysis of key biological processes, cellular compartments and molecular function associated with 797 differentially expressed genes between neural crest iPericytes and mesoderm iPericytes.

Next, we assessed for differential gene expression between mesoderm and neural crest iPericytes, which revealed 458 genes upregulated and 339 genes downregulated in neural crest iPericytes compared to mesoderm iPericytes (Fig. 2E). Visualisation of these differentially expressed genes via a heat map demonstrated the separation between mesoderm iPericytes and neural crest iPericytes (Fig. 2F). Gene ontology analysis showed enrichment for genes related to tissue development, extracellular matrix production, DNA/RNA processing and growth factor binding and activity (Fig. 2G). These differences could reflect changes in cellular function between mesoderm and neural crest iPericytes and HBVPs.

### Validation of the mesoderm iPericyte differentiation protocol using multiple iPSC lines

To confirm that iPericyte differentiation is highly reproducible, multiple unrelated iPSC lines (MNZTASi019-A, MNZTASi021-A, and MNZTASi022-A) were cultured and used to generate mesoderm iPericytes. RNA was collected from the iPSCs and the iPericytes for bulk RNA sequencing. PCA of the gene expression profile of the iPSCs and mesoderm iPericytes revealed that each cell type (iPSCs and iPericytes) clustered separately along the first principal component, accounting for 93% of sample variation (Fig. 3A). Variation between replicates accounted for only 5% of sample variation, showing a remarkable similarity between replicates (Fig. 3A). We then selected genes associated with iPSC, pericyte, endothelial cell, microglia, oligodendrocyte progenitor cell (OPC), oligodendrocyte, astrocyte, or neuronal identity, and generated a heat map of gene expression for each iPSC line and the corresponding iPericytes (Fig. 3B). Regardless of donor, iPericytes had successfully downregulated the pluripotency genes *NANOG*, *POU5F1* and *SOX2*, and upregulated pericyte-associated genes, including *PDGFRB, CSPG4, ANPEP* and *ACTA2* (Fig. 3B). Gene expression was consistent across iPericytes generated from different iPSC lines (Fig. 3B). Importantly, iPericytes did not express genes synonymous with other neurovascular cell types (Fig. 3B). These data indicate this differentiation protocol can be applied to distinct iPSC lines and produce iPericytes with a consistent mRNA expression profile.

**Figure 3.**
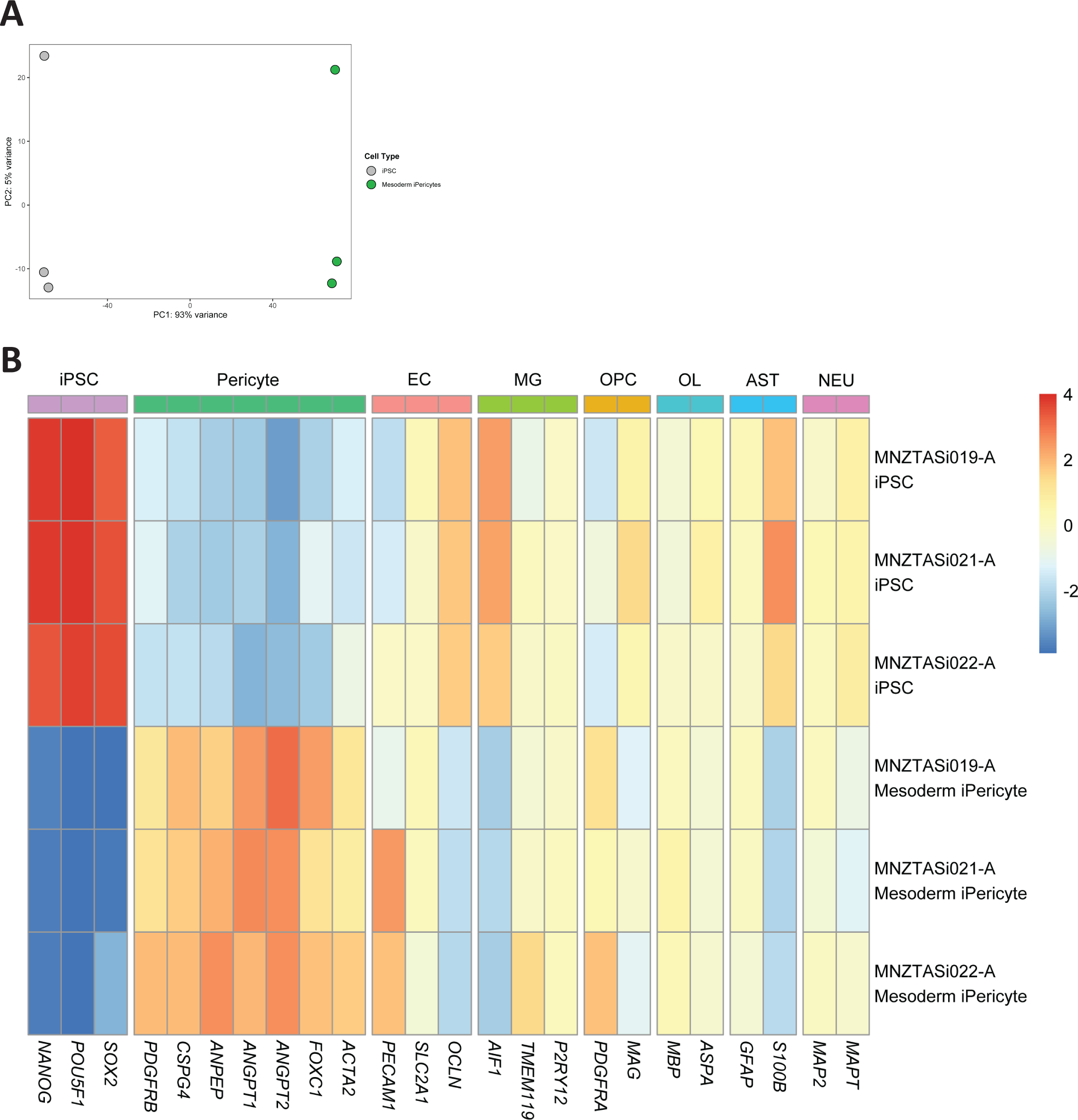
Mesoderm iPericytes from multiple cell lines have similar mRNA expression. (A) Principal components analysis showing separate clustering of mesoderm iPericytes and iPSCs from n = 3 different cell lines. (B) Heat map showing relative expression levels in iPSCs and mesoderm iPericytes of key genes typically expressed by iPSCs, pericytes, endothelial cells (EC), microglia (MG), oligodendrocyte precursor cells (OPCs), oligodendrocytes (OL), astrocytes (AST) and neurons (NEU). Warmer colours indicate higher expression, cooler colours indicate lower expression.

### PDGFRβ signalling promotes iPericyte proliferation

mRNA expression differences between HBVPs and iPericytes could influence their capacity to respond to environmental signals, and so we next compared the proliferative capacity of these cells. A key ligand-receptor pathway that pericytes utilise for survival and proliferation is the PDGFRβ signalling pathway (20). We exposed HBVPs or iPericytes to basal pericyte medium alone (PM) or PM containing the PDGFRβ ligand, PDGF-BB (100 ng/ml), in the presence of the thymidine analogue, EdU, as previously described (20). The addition of PDGF-BB increased the proportion of HBVPs and iPericytes that incorporated EdU over a 24 h period, indicative of increased proliferation (Fig. 4A &B, Fig.S3; HBVP, p < 0.0001; neural crest iPericytes, p = 0.01; mesoderm iPericytes, p < 0.0001). The magnitude of response to PDGF-BB was similar between all three pericyte lines. Similar results were observed when complete pericyte media (CPM), containing specialised pericyte growth supplement (ScienCell, USA), was used compared to PM (Fig. 4B). These results indicate that iPericytes can proliferate in response to the pericyte growth factor PDGF-BB.

**Figure 4.**
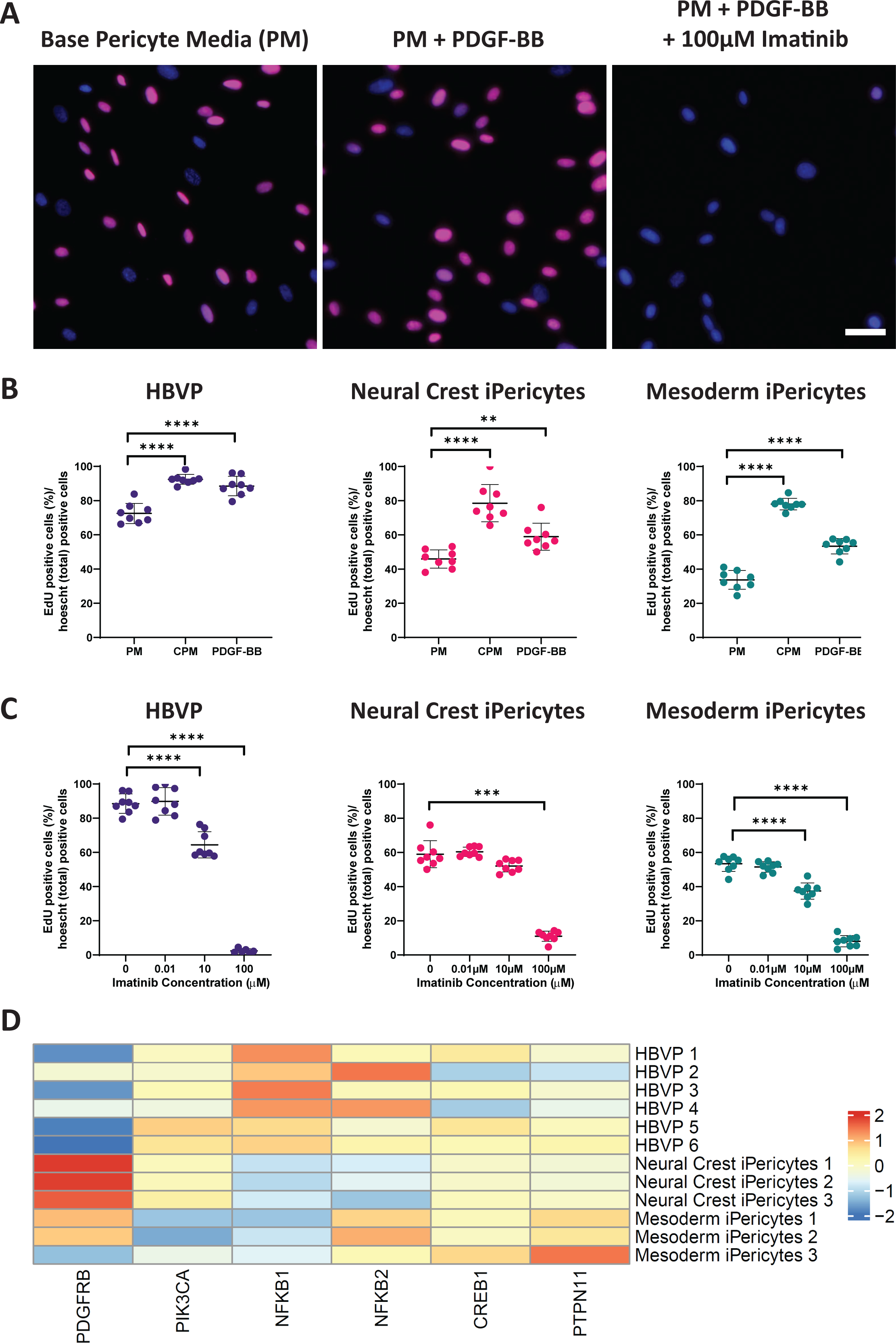
Proliferation of iPericytes through the PDGF-BB: PDGFRβ signalling pathway. (A) iPericytes were incubated in basal pericyte media (PM) and treated with PDGF-BB (PM + PDGF-BB) while being exposed to 100 µM imatinib (PM + PDGF-BB + 100µM imatinib). Proliferation was measured using an EdU uptake assay. iPericytes that are EdU-positive are indicated by magenta, while total number of iPericytes were measured by DAPI (blue). Scale bar = 50 µm. (B) Quantification of HBVPs, neural crest iPericytes and mesoderm iPericytes proliferating (as indicated by EdU-positive staining) as a percentage of total cells following 24 h exposure to PM, complete pericyte media with pericyte growth factors (CPM) or PM + PDGF-BB (n = 8 per condition). Data were analysed using a one-way ANOVA: HBVP (F (2, 21) = 35.52, p < 0.0001); neural crest iPericyte (F (2, 21) = 30.85, p < 0.0001); mesoderm iPericyte (F (2, 21) = 191.4, p < 0.0001). (C) Quantification of changes to PDGF-BB-induced proliferation with increasing concentrations of imatinib over 24 h in HBVPs, neural crest iPericytes and mesoderm iPericytes (n = 8 per condition). Data were analysed using a one-way ANOVA or Kruskal-Wallis test: HBVP (F (3, 26) = 259.2, p < 0.0001); neural crest iPericyte (H (3) = 24.41, p < 0.0001); mesoderm iPericyte (F (3, 28) = 221.5, p < 0.0001). For (B) and (C), post-hoc comparisons were performed using Dunnett’s multiple comparisons or Dunn’s test: * p < 0.05, ** p < 0.01, *** p < 0.001,**** p < 0.0001. Data shown as mean ± SD. **(D)** Heat map of key genes involved in pericyte proliferation in the PDGF-BB: PDGFRβ signalling pathway in HBVP, neural crest iPericytes and mesoderm iPericytes selected from Sweeney et al. 2016 (27).

To confirm that the proliferative response was mediated by PDGFRβ, HBVP and iPericyte proliferation was assessed in the presence of imatinib. In pericytes, imatinib inhibits PDGFRβ phosphorylation to prevent proliferation (20). In HBVPs and iPericytes, imatinib produced a dose dependent inhibition of PDGF-BB-induced proliferation (Fig. 4C, Fig. S3). For HBVPs, 0.01 µM imatinib did not alter proliferation (p= 0.9851), while 10 µM imatinib and 100 µM imatinib significantly reduced proliferation by 31% and 96% of PDGF-BB alone, respectively (p < 0.0001). Mesoderm iPericytes also failed to respond to 0.01µM imatinib (51%, p = 0.6517), while 10 µM and 100 µM imatinib significantly reduced proliferation to 37% and 8% of PDGF-BB alone, respectively (p < 0.0001). Neural crest iPericytes were less sensitive to PDGFRβ blockade, as neither 0.01µM (p = 0.9723) or 10 µM (p = 0.3121) altered PDGF-BB-induced proliferation. However, 100 µM imatinib significantly reduced the proliferation rate to 11% of that recorded for PDGF-BB alone (p = 0.0009). These findings indicate that neural crest iPericytes are less sensitive than mesoderm iPericytes or HBVPs to PDGFRβ inhibition.

To determine why neural crest iPericytes have altered susceptibility to PDGFRβ inhibition, we interrogated our RNA-sequencing dataset, and identified differences between HBVPs, mesoderm and neural crest iPericytes, in the relative expression of mRNAs downstream of the PDGF-BB:PDGFRβ pathway. In particular, *PIK3CA* (log2FoldChange = −0.67, p_adj_ = 2.76E^-5^), *NFKB1* (log2FoldChange = −1.28, p_adj_ = 2.47E^-26^), *NFKB2* (log2FoldChange = −0.78, p_adj_ = 0.00096), *CREB1* (log2FoldChange = −0.46, p_adj_ = 2.39E^-6^) and *PTPN11* (log2FoldChange = −0.36, p_adj_ = 0.003) were differentially expressed between HBVPs and iPericytes, while *PIK3CA* (log2FoldChange = −0.44, p_adj_ = 0.039) and *NFKB2* (log2FoldChange = 0.68, p_adj_ = 2.24E^-5^) were differentially expressed between mesoderm and neural crest iPericytes (Fig. 4D). RNAseq analysis also revealed that the expression of *PDGFRβ* was significantly higher (log2FoldChange = −1.06, p_adj_ = 6.26E^-6^) in neural crest iPericytes compared to mesoderm iPericytes (Fig. 4D), which is in line with the qPCR data (Fig. 1B). These differences could explain why neural crest iPericytes required a higher concentration of imatinib to prevent PDGF-BB mediated proliferation.

### iPericytes contract in response to endothelin-1

A primary function of pericytes is to contract and dilate to modulate capillary diameter, thereby altering cerebral blood flow (4). We previously used a single cell imaging assay (23) and the xCelligence electrical impedance assay (22) to show that HBVPs can respond to vasoactive mediators. To assess the responses of mesoderm and neural crest iPericytes to endothelin-1, we again used the xCelligence system. Cells were plated on specialised cell culture plates that allow resistance to electron flow to be measured to provide an assessment of cell index (Fig. S4A). Normalised cell index values can be analysed to compare differences in slope, AUC and change in cell area after treatment with contractile mediators (Fig. S4B). It is important to note that a small reduction in normalised cell index is ordinarily observed over the first few minutes of an experiment, even under control conditions (Fig. 5A & B, (22)). When mesoderm iPericytes (Fig. 5A) and neural crest iPericytes (Fig. 5B) were treated with endothelin-1, normalised cell index decreased compared to vehicle suggesting pericytes had contracted, which was confirmed when AUC was calculated (treatment: p = 0.0033, Fig. 5C; treatment: p < 0.0001, Fig. 5F). Compared to HBVPs, contraction of mesoderm iPericytes (p = 0.9995, Fig. 5C) and neural crest iPericytes (p = 0.1464, Fig. 5F) was similar in the first 20 min of endothelin-1 exposure. The maximum contraction achieved by mesoderm iPericytes was the same as HBVPs in response to endothelin-1 (treatment: p = 0.0021, Fig. 5D), and this was maintained over 2 h (treatment: p = 0.0026, Fig. 5E). However, there was a different effect of treatment with endothelin-1 on neural crest iPericytes in comparison to HBVPs (interaction of cell type x treatment: p = 0.0010, Fig. 5G). Post-hoc analysis revealed that neural crest iPericytes maximum contraction was greater in response to endothelin-1 compared to HBVPs (p = 0.0007, Fig. 5G) and they also sustained a greater level of contraction compared to HBVPs for up to 2 h (p = 0.0001, Fig. 5H). These findings suggest that iPericytes derived through different lineages display distinct responses to endothelin-1.

**Figure 5.**
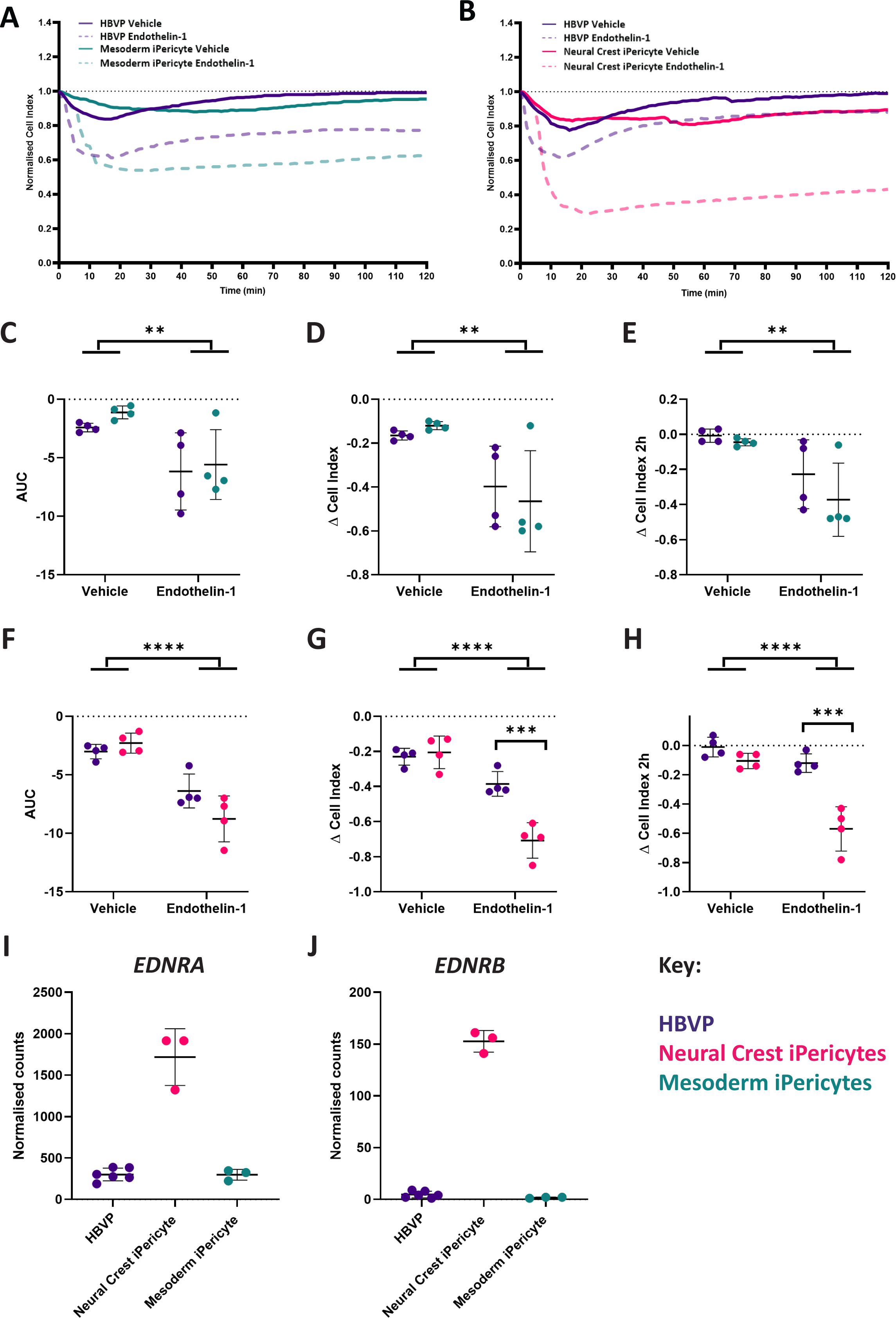
Endothelin-1 induces iPericyte contraction. (A-B) Normalised cell index of neural crest iPericytes, mesoderm iPericytes and HBVPs treated with endothelin-1 or vehicle (CPM) over a period of 2 h (n = 4 per condition). (C-E) Quantified AUC, Δ cell index and Δ cell index after 2 h for mesoderm iPericytes and HBVPs treated with control or endothelin-1 analysed using two-way ANOVA: AUC (cell type: F (1, 12) = 0.6953, p = 0.4206; treatment: (F (1, 12) = 13.35, p =0.0033; interaction: F (1, 12) = 0.1006, p = 0.7565); Δ cell index (cell type: F (1, 12) = 0.02309, p = 0.8817; treatment: F (1, 12) = 15.21, p = 0.0021; interaction: F (1, 12) = 0.5773, p = 0.4620); Δ cell index after 2 h (cell type: F (1, 12) = 1.590, p = 0.2313; treatment: F (1, 12) = 14.31, p = 0.0026; interaction: F (1, 12) = 0.5518, p = 0.4719). (F-H) Quantified AUC (indicator of volume of contraction), Δ cell index (maximum contraction) and Δ cell index after 2 h (contraction at 2 h time point) for neural crest iPericytes and HBVPs treated with control or endothelin-1 analysed using two-way ANOVA: AUC (cell type: F (1, 12) = 1.563, p = 0.2351; treatment: (F (1, 12) = 54.67, p < 0.0001; interaction: F (1, 12) = 5.470, p = 0.0375); Δ cell index (cell type: F (1, 12) = 13.53, p = 0.0032; treatment: F (1, 12) = 66.11, p < 0.0001; interaction: F (1, 12) = 18.47, p = 0.0010); Δ cell index after 2 h (cell type: F (1, 12) = 34.64, p < 0.0001; treatment: F (1, 12) = 38.56, p < 0.0001; interaction F (1, 12) = 14.70, p = 0.0024). (C-H) Post-hoc comparisons performed using Sidak’s multiple comparisons test. * p < 0.05, ** p < 0.01, *** p < 0.001,**** p < 0.0001. Data shown as mean ± SD. (I-J) Normalised gene expression counts of differentially expressed endothelin-1 receptors in HBVP, neural crest iPericytes and mesoderm iPericytes compared using DEseq: HBVPs and neural crest iPericytes *EDNRA* Fig. 5I log2FoldChange = 2.53, p_adj_ = 6.12E^-23^; *EDNRB* Fig. 5J log2FoldChange = 5.14, p_adj_ = 9.13E^-26^; neural crest iPericytes compared to mesoderm iPericytes *EDNRA* (I) log2FoldChange = −2.52, p_adj_ = 3.12387E^-25^; *EDNRB* (J) log2FoldChange = −7.22, p_adj_ = 6.77016E^-15^.

To determine whether the lineage specific responses of iPericytes were due to differences in endothelin-1 receptor expression, we determined whether endothelin-1 receptor genes were differentially expressed between HBVPs, neural crest iPericytes and mesoderm iPericytes. *EDNRA* and *EDNRB*, genes which code for the two major endothelin-1 receptors, were differentially expressed in our RNA-seq dataset. There was a significantly different expression of both subtypes of endothelin-1 receptor between HBVPs and neural crest iPericytes (*EDNRA* Fig. 5I log2FoldChange = 2.53, p_adj_ = 6.12E^-23^; *EDNRB* Fig. 5J log2FoldChange = 5.14, p_adj_ = 9.13E^-26^), while there was no difference between HBVP and mesoderm iPericytes (Fig. 5I, J). There was also significantly higher expression of both subtypes of endothelin-1 receptor in neural crest iPericytes compared to mesoderm iPericytes (*EDNRA* Fig. 5I log2FoldChange = −2.52, p_adj_ = 3.12E^-25^; *EDNRB* Fig. 5J log2FoldChange = −7.22, p_adj_ = 6.77E^-15^), which might be driving their greater response to the endothelin-1 ligand. These data indicate that iPericytes can respond to endothelin-1, and that neural crest iPericytes display a greater contractile response to endothelin-1 compared to mesoderm iPericytes and HBVPs.

### iPericytes have functional responses to the vasodilator adenosine

Given we observed differences in the response of neural crest iPericytes and mesoderm iPericytes to endothelin-1, we also tested the response of iPericytes to adenosine which can initiate pericyte relaxation *in vitro* (22). Similar to HBVPs, when mesoderm iPericytes (Fig. 6A) and neural crest iPericytes (Fig. 6B) were exposed to adenosine, normalised cell index increased compared to vehicle conditions, indicative of pericyte relaxation. When treated with adenosine, mesoderm iPericytes relaxed (treatment: p = 0.0002, Fig. 6C) and the maximum relaxation achieved by mesoderm pericytes was the same as HBVPs in response to adenosine (treatment: p = 0.0002, Fig. 6D), however, this was not maintained over 2 h (treatment: p = 0.7317, Fig. 6E). There was a different response following adenosine treatment on neural crest iPericyte relaxation in comparison to HBVPs (interaction of cell type x treatment: p = 0.0202, Fig. 6F). Post-hoc analysis revealed that neural crest iPericytes relax less in response to adenosine compared to HBVPs (p = 0.0112, Fig. 6F), which was also observed in assessment of maximum relaxation (p = 0.0336, Fig. 6G) and relaxation at 2 h (p = 0.0170, Fig. 6H). These findings indicate that neural crest iPericytes display reduced ability to relax in response to adenosine compared to mesoderm iPericytes.

**Figure 6.**
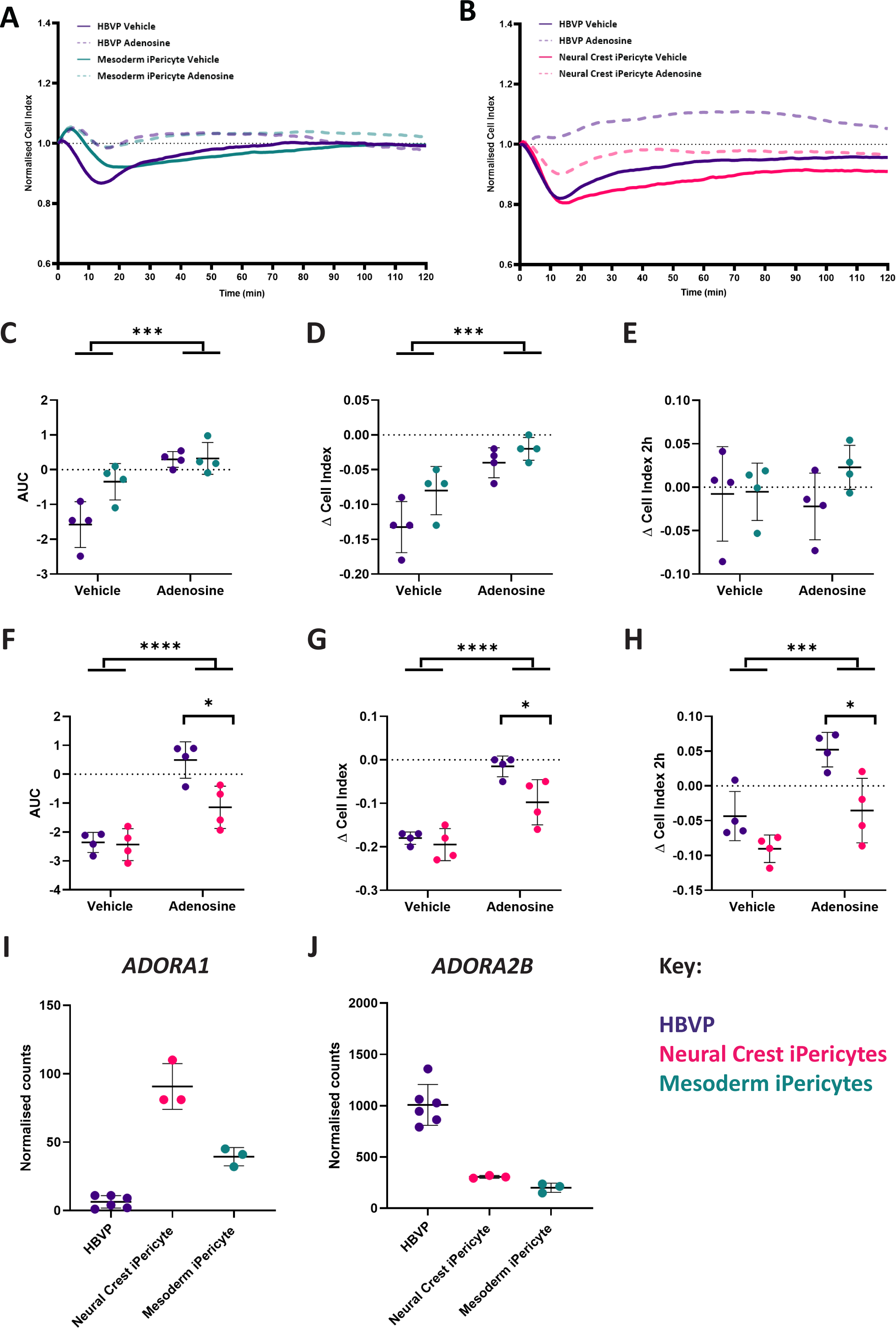
Adenosine induces iPericyte relaxation. (A-B) Normalised cell index of neural crest iPericytes, mesoderm iPericytes and HBVPs treated with adenosine or vehicle (CPM) over a period of 2 h (n = 4 per condition). (C-E) Quantified AUC, Δ cell index and Δ cell index after 2 h for mesoderm iPericytes and HBVPs treated with control or adenosine analysed using two-way ANOVA: AUC (cell type: F (1, 12) = 6.583, p = 0.0247; treatment: (F (1, 12) = 26.84, p =0.0002; interaction: F (1, 12) = 6.027, p = 0.0303); Δ cell index (cell type: F (1, 12) = 6.387, p = 0.0265; treatment: F (1, 12) = 28.26, p = 0.0002; interaction: F (1, 12) = 1.284, p = 0.2794); Δ cell index after 2 h (cell type: F (1, 12) = 1.460, p = 0.2502; treatment: F (1, 12) = 0.1232, p = 0.7317; interaction: F (1, 12) = 1.174, p = 0.2999). (F-H) Quantified AUC, Δ cell index and Δ cell index after 2 h for neural crest iPericytes and HBVPs treated with control or adenosine analysed using two-way ANOVA: AUC (cell type: F (1, 12) = 8.596, p = 0.0126; treatment: (F (1, 12) = 50.38, p < 0.0001; interaction: F (1, 12) = 7.159, p = 0.0202); Δ cell index (cell type: F (1, 12) = 7.881, p = 0.0158; treatment: F (1, 12) = 57.12, p < 0.0001; interaction: F (1, 12) = 3.777, p = 0.0758); Δ cell index after 2 h (cell type: F (1, 12) = 16.46, p = 0.0016; treatment: F (1, 12) = 20.58, p = 0.0007; interaction: F (1, 12) = 1.500, p = 0.2442). (C-H) Post-hoc comparisons performed using Sidak’s multiple comparisons test. * p < 0.05, ** p < 0.01, *** p < 0.001,**** p < 0.0001. Data shown as mean ± SD. (I-J) Normalised gene expression counts of differentially expressed adenosine receptors in HBVP, neural crest iPericytes and mesoderm iPericytes compared using DEseq: HBVPs and neural crest iPericytes *ADORA1* (I) log2FoldChange = 3.92, p_adj_ = 1.65E^-11^; *ADORA2B* (J) log2FoldChange = −1.71, p_adj_ = 2.93E^-19^; HBVPs and mesoderm iPericytes *ADORA1* (I) log2FoldChange = 2.72, p_adj_ = 0.00006; *ADORA2B* (J) log2FoldChange = −2.33, p_adj_ = 1.52E^-25^; neural crest iPericytes compared to mesoderm iPericytes *ADORA1* (I) log2FoldChange = −1.19, p_adj_ = 0.001; *ADORA2B* (J) log2FoldChange = −0.61, p_adj_ = 0.03288.

To determine whether the lineage specific responses of iPericytes were due to differences in adenosine receptor expression, we determined whether adenosine receptor genes were differentially expressed between HBVPs, neural crest iPericytes and mesoderm iPericytes. *ADORA1* and *ADORA2B*, genes which code for two of the major adenosine receptors, were differentially expressed in our RNA-seq dataset. There was a significantly different expression of adenosine receptors type *ADORA1* and *ADORA2B* between HBVPs and neural crest iPericytes (*ADORA1* Fig. 6I log2FoldChange = 3.92, p_adj_ = 1.65E^-11^; *ADORA2B* Fig. 6J log2FoldChange = −1.71, p_adj_ = 2.93E^-19^). There was also significantly higher expression of both of these subtypes in neural crest iPericytes compared to mesoderm iPericytes (*ADORA1* Fig. 6I log2FoldChange = −1.19, p_adj_ = 0.001; *ADORA2B* Fig. 6J log2FoldChange = −-0.61, p_adj_ = 0.033). Therefore, while differences in the expression of adenosine receptors exist between HBVPs, mesoderm and neural crest iPericytes, they do not reflect differences in functional responses to adenosine. These findings suggest that iPericytes can respond to adenosine, and that neural crest iPericytes display a reduced relaxation response compared to HBVPs.

## Discussion

iPSC-derived neurovascular cells are becoming a popular model of choice to investigate the function of the NVU *in vitro*. Methods to generate iPericytes reveal a novel avenue that will allow researchers to determine how patient specific genetic variants affect pericyte function, will help to create more accurate *in vitro* models of the NVU and disease and, ultimately, may provide a reproducible and personalised tool for implantation in regenerative medicine. To confidently use these cells to study pericyte function, it is important to establish how representative they are of primary pericytes *in vitro*. We derived mesoderm and neural crest iPericytes using a previously published protocol (16) and showed they express classical pericyte mRNAs, but do not express other brain cell markers. We then found that there were differences between mesoderm and neural crest iPericytes in their functional response to the PDGF-BB:PDGFRβ signalling pathway that mediates proliferation, and in response to known vasoactive mediators endothelin-1 and adenosine, in comparison to HBVPs.

### Mesoderm and neural crest iPericytes express key pericyte markers and morphologies

Using both qPCR and immunocytochemistry, we sought to test the expression of key pericyte mRNAs or proteins in neural crest and mesoderm iPericytes derived from the TOB-00220 line. We found that both developmental lineages of iPericytes express three classical pericyte mRNAs *ANPEP* (encoding CD13), *CSPG4* (encoding NG2) and *PDGFRB* (encoding PDGFRβ), with immunocytochemistry confirming their protein expression, in line with a previous study (16). Importantly, we also compared iPericyte mRNAs expression to primary HBVPs and showed high levels of expression compared to iPSCs. iPericytes downregulate expression of key pluripotency mRNAs *OCT4* and *NANOG,* showing a distinct change compared to the iPSCs from which they were derived. In addition, iPericytes display the five morphological subtypes previously described for primary HBVPs (23). The majority of cells exhibited standard morphology, which we have previously shown possess contractile capacity (23). These data highlight that iPericytes express pericyte markers and are morphologically similar to HBVPs.

### iPericytes retain lineage specific differences in gene expression

We next sought to understand whether separate iPericyte lineages could display altered gene expression and which biological processes these were related to. Following RNA sequencing, we showed that gene expression in HBVPs was markedly different compared to both mesoderm and neural crest iPericytes. Differentially expressed genes appeared to be related to tissue development and protein binding, which could impact the function of these cells. It has already been shown that pericytes contribute to the development of the vascular network in multiple organs and they can assist in the development of key cellular structures such as the extracellular matrix (25). It is possible that differences in the genetic background of HBVPs compared to the iPSC line we used could explain the extent of differential gene expression. In addition, HBVPs could include both mesoderm and neural crest-derived pericytes given that both lineages reside in the brain (7). We also identified that there were gene expression differences between neural crest and mesoderm iPericytes that was related primarily to organ development. These differences could be explained by the mesoderm lineage being more prominent in organ development throughout the body whereas cells from the neural crest pathway would be restricted to the nervous system (26). Interestingly, genes related to growth factor binding and activity were also differentially expressed which could indicate differences in function of pericytes derived from these two lineages. This was evident in both the proliferation and contractility assays where functional responses differed, suggesting pericytes of different lineages may have altered physiological responses.

### iPericytes can be consistently produced from different iPSC lines

Using RNA sequencing, we showed that gene expression profiles of iPericytes that had been differentiated from three separate iPSC lines derived from three unrelated individuals were consistent between different lines. In particular, all three iPericyte lines had consistent levels of enriched pericyte mRNA expression, while downregulating expression of known stem cell genes. Notably, iPericytes did not express key markers of any other cell type such as endothelial cells, microglia, OPCs, oligodendrocytes, astrocytes, or neurons. This suggests that iPericytes can be produced with high consistency from different iPSC lines, supporting their use for assessing pericyte function in disease contexts.

### iPericytes proliferate in response to the PDGFRβ ligand PDGF-BB

Although consistent expression of key pericyte mRNAs and proteins by iPericytes is encouraging, it is important that this translates into functional characteristics representative of pericytes *in vivo*. To expand our knowledge on the relative similarities between mesoderm and neural crest iPericytes, we compared their functional response to PDGF-BB, a growth factor essential for pericyte proliferation and survival (27). We have previously shown that HBVPs proliferate in response to PDGF-BB *in vitro* through the PDGFRβ receptor (20). Like HBVPs, mesoderm and neural crest iPericytes proliferated in the presence of PDGF-BB. In addition, the specificity of this proliferative response to the PDGFRβ pathway was confirmed by the blockade of this response with the PDGFRβ inhibitor imatinib, similar to HBVPs. The PDGF-BB:PDGFRβ signalling pathway is essential for pericyte and endothelial cell interactions at the NVU, mediating key endothelial cell processes such as angiogenesis (27). A number of studies have assessed iPericytes in co-culture with endothelial cells (8–10, 15), showing that iPericytes can specifically support endothelial tube formation and the strength of the endothelial barrier through trans-endothelial resistance measures (7–10, 16). In addition, iPericytes have been used as part of functional blood-brain barrier models (7, 28–30). Furthermore, a recent study showed the capacity of iPericytes to aid in BBB repair in pericyte deficient mice, suggesting functional signalling between endothelial cells and iPericytes is also possible *in vivo* (31). These studies highlight the capacity for iPericytes to support and enhance survival and differentiation of other key cells of the NVU.

Until now functional studies of iPericytes have typically focussed on one pericyte developmental lineage at a time, either mesoderm (8–10) or neural crest (7, 11), restricting comparisons between the two. Here, we demonstrate for the first time that neural crest iPericytes display altered PDGFRβ signalling responses compared to HBVPs and mesoderm iPericytes, with a higher concentration of the PDGFRβ receptor inhibitor imatinib required to inhibit proliferation *in vitro*. This finding was supported by higher expression of the *PDGFRB* gene by neural crest iPericytes. These differences may reflect an inherent difference in the function of this receptor pathway between neural crest and mesoderm pericytes that should be considered for future studies.

### iPericytes are responsive to the vasoactive mediators endothelin-1 and adenosine

Another key function of pericytes is their role in blood flow regulation. It has previously been shown that pericytes possess the contractile protein αSMA which can generate a contractile response in these cells (32, 33). However, there has been some discordance in the literature about expression of αSMA and contractility of pericytes (4, 34). This discordance has also been observed with iPericytes *in vitro* with some studies showing αSMA expression in iPericytes (10, 11) and some concluding that it is not expressed (7, 8). Interestingly, Kumar *et al*. (10) found that αSMA expression could be triggered through a specific pericyte differentiation protocol involving PDGF-BB, vascular endothelial growth factor (VEGF), activin receptor-like kinase receptor (ALK) inhibitor SB-431542 and epidermal growth factor (EGF), suggesting certain growth factors must be present for expression of αSMA in iPericytes. Here, we showed that the αSMA gene *ACTA2* was expressed in both neural crest and mesoderm iPericytes, with bulk RNA sequencing revealing reproducible expression of *ACTA2* in mesoderm iPericytes throughout three separate iPSC lines. Given that HBVP expression of αSMA was associated with contractile ability (23), the expression of αSMA is suggestive of the potential to contract.

It has previously been shown that HBVPs contract in response to endothelin-1 and relax in response to adenosine *in vitro* (22). Using a similar approach, we found that exposing neural crest and mesoderm iPericytes to endothelin-1 led to a strong reduction in cell area indicative of cell contraction. Interestingly, neural crest iPericytes had a much stronger contractile response to endothelin-1 compared to both HBVPs and mesoderm iPericytes. Further analysis into gene expression changes revealed that the two major endothelin-1 receptors (*EDNRA* and *EDNRB*) were more highly expressed in neural crest iPericytes. In addition, the expression of *ACTA2*, the gene encoding the key contractile protein αSMA, was more highly expressed by neural crest iPericytes. However, neural crest iPericytes appear to not relax in response to adenosine as much as HBVPs and mesoderm iPericytes. Given that neural crest iPericytes expressed similar (*ADORA2B*) or higher (*ADORA1*) levels of adenosine receptor compared to mesoderm iPericytes, this suggests other factors may be influencing the extent to which neural crest iPericytes react to adenosine. Overall, these experiments highlight the ability of iPericytes to contract and relax, with some reactivity differences between lineages.

## Conclusion

Collectively, we illustrate that neural crest and mesoderm iPericytes, derived from multiple iPSC lines, are morphologically similar to HBVPs and express key pericyte markers. iPericytes are functionally active, demonstrated through proliferation in response to the key pericyte growth factor PDGF-BB, contraction in response to endothelin-1, and relaxation in response to adenosine. These findings suggest that iPericytes behave functionally like HBVPs, providing further support for their use as a tool to study pericyte function. We observed some differences between iPericytes of different lineages, notably that neural crest iPericytes were less sensitive to PDGFRβ inhibition and more contractile compared to mesoderm iPericytes. Therefore, differences between iPericytes derived through different lineages must be taken into consideration when designing experiments using iPericytes to assess pericyte function.

## Author Contributions

NEK: methodology; validation; investigation; formal analysis; visualisation; writing - original draft; writing – review and editing. J-MC: methodology; software; formal analysis; investigation; resources; data curation; writing-review and editing; visualisation; supervision. LSB: methodology; formal analysis; investigation; writing-review and editing; visualisation. AJF: investigation; methodology; formal analysis; visualisation; writing – review and editing. NBB: data curation; methodology; formal analysis; visualisation; supervision; writing – review and editing. JLF: investigation; methodology; supervision; writing - review and editing. JMC: methodology; investigation; writing-review and editing. JT: methodology; writing – review and editing. AP: methodology; resources; writing – review and editing. AWH: methodology; resources; writing – review and editing. GPM: methodology; supervision; writing - original draft; writing – review and editing. KMY: conceptualisation; funding acquisition; methodology; project administration; resources; supervision; writing – review and editing. ALC: conceptualisation; funding acquisition; methodology; project administration; resources; supervision; writing – review and editing. BAS: conceptualisation; funding acquisition; methodology; project administration; resources; supervision; writing – original draft; writing – review and editing.

## Acknowledgements

We would like to thank study participants who donated cells for iPSC generation. The University of Tasmania provided core laboratory facilities for this work. We would like to acknowledge the use of the high-performance computing facilities provided by Digital Research at the University of Tasmania. We would like to acknowledge the Australian Genome Research Facility (AGRF) who performed all of the RNA sequencing for gene expression analysis.

## Data Availability

Please contact Kaylene Young to source MS Stem iPSCs. Imaging data are available from the corresponding author upon reasonable request. Bulk RNA sequencing data will be made available online upon a revised submission.

