## Supplementary material for "Induced pluripotent stem cell derived pericytes respond to endogenous mediators of proliferation and contractility"

*Corresponding author:

### Supplementary Methods

#### Pericyte morphology analysis

Phase contrast micrographs of neural crest and mesoderm iPericytes were captured with the 4x objective on a Zeiss microscope using a Camera AxioCam ICc5 (Imbros). Each individual pericyte was manually assigned to a morphological subtype based on their appearance according to the criteria outlined previously (1). Percentage of each morphology within cultures was calculated from N = 746 (neural crest iPericytes) and 759 (mesoderm iPericytes) cells analysed across one culture for each differentiation method.

#### RNA Sequencing and differential gene expression analysis

*Comparison of neural crest iPericytes to mesoderm iPericytes*

Following library preparation for each sample, RNA sequencing (100bp single-end run) was carried out using an Illumina NovaSeq platform. Following demultiplexing and quality control, the cleaned sequence reads were aligned against the *Homo sapiens* genome (build version hg38) using the STAR aligner (v2.7.10a) (9). Raw gene counts were generated and transcripts were assembled with the StringTie tool (v2.1.4) (10). Differential gene expression analysis was conducted using DESeq2 (v1.36.0) (11) in R (v4.2.1) with an adjusted p-value (false discovery rate) $<0.05$ and absolute log2(fold-change) $> 1$ considered to be differentially expressed. Gene ontology enrichment analysis was conducted using clusterProfiler (v4.4.4) (12).

*Comparison of three iPSC lines differentiated to three mesoderm iPericyte lines*

Following library preparation for each sample, RNA was sequenced to a minimum depth of 20 million reads with 150 base paired end lengths. Raw sequencing data was processed with TrimGalore (v0.6.7) (2). The sequencing adapter (CTGTCTCTTATACACATCT) was trimmed from each read and 1 base pair was removed from the 5' end of each read. A read quality Phred score threshold of 30 was used to remove low quality reads. Reads with a length of less than 25 base pairs after trimming were discarded. The quality of the sequencing data was then evaluated using FastQC (v.0.11.9) (3), and all data met the necessary quality metrics for analysis.

Gene expression was quantified using Salmon (v.1.8.0)(4). The Salmon indexed transcriptome for *Homo sapiens* (GRCh38) was downloaded using RefGenie (v.0.12.1) (5) and Salmon used to quantify gene expression against this transcriptome in mapping-based mode. The sequencing library type was ISR. Within Salmon, --validateMappings was employed to improve the sensitivity and specificity of the read mapping (and thus quantification accuracy); --seqBias and --gcBias were employed to enable Salmon to learn and correct for sequence-specific biases and fragment-level GC biases respectively. The result of this analysis was per sample quantification of transcript expression.

Differential gene expression analysis was performed using DESeq2 (v1.34.0) (6) to compare gene expression between iPSCs and mesoderm-iPericytes. Tximport (v1.27.1) (7) was used to import transcript-level abundance into R (v.4.3.0) and summarise abundance to the gene-level. Genes with less than 1 read across each group were filtered out. Heatmaps were generated using pheatmap (v.1.0.12) (8).

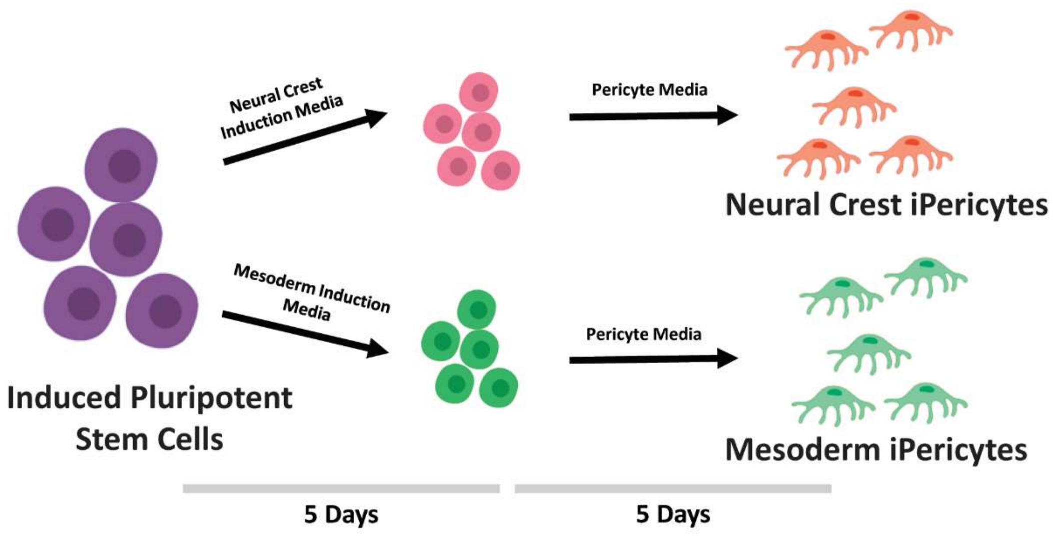


**Supplementary Figure 1. iPSCs can be differentiated into pericytes through the neural crest and mesoderm induction pathways.** iPSCs were placed in either mesoderm induction media or neural crest induction media for 5 days. Media was then replaced with complete pericyte media for another 5 days to generate neural crest iPericytes or mesoderm iPericytes. Protocol adapted from Faal et al. (13).

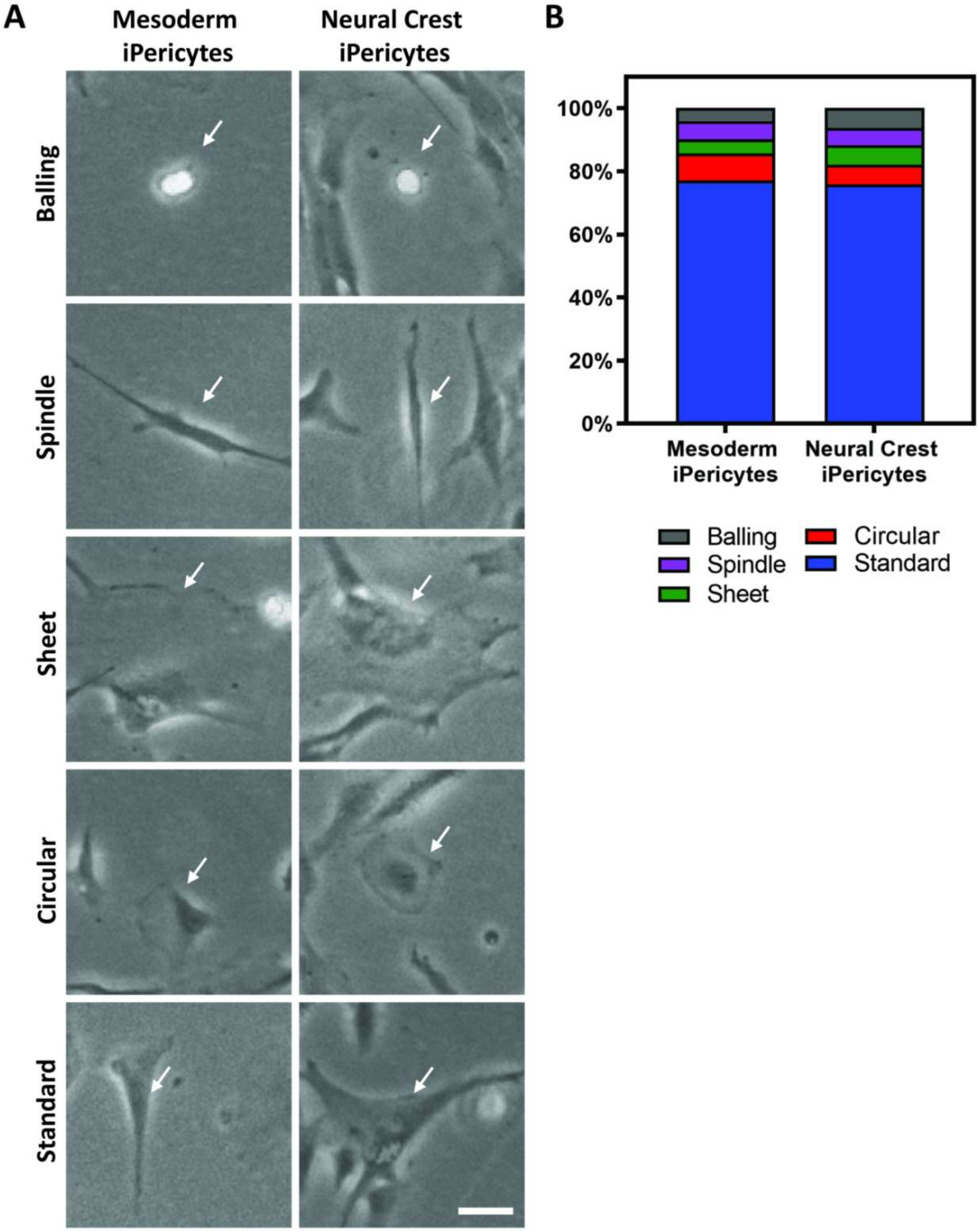


**Supplementary Figure 2. iPericytes display heterogenous morphology consistent with HBVP cells *in vitro*.** Pericyte morphology was classified following the criteria outlined in Brown et al. (1). (A) Both neural crest iPericytes and mesoderm iPericytes display the five key morphological classifications of pericytes *in vitro*: balling, spindle, sheet, circular, and standard. (B) Quantification of the percentage of each pericyte morphology subtype in neural crest iPericyte culture (746 cells) and mesoderm iPericyte culture (759 cells). Scale = 20 µm.

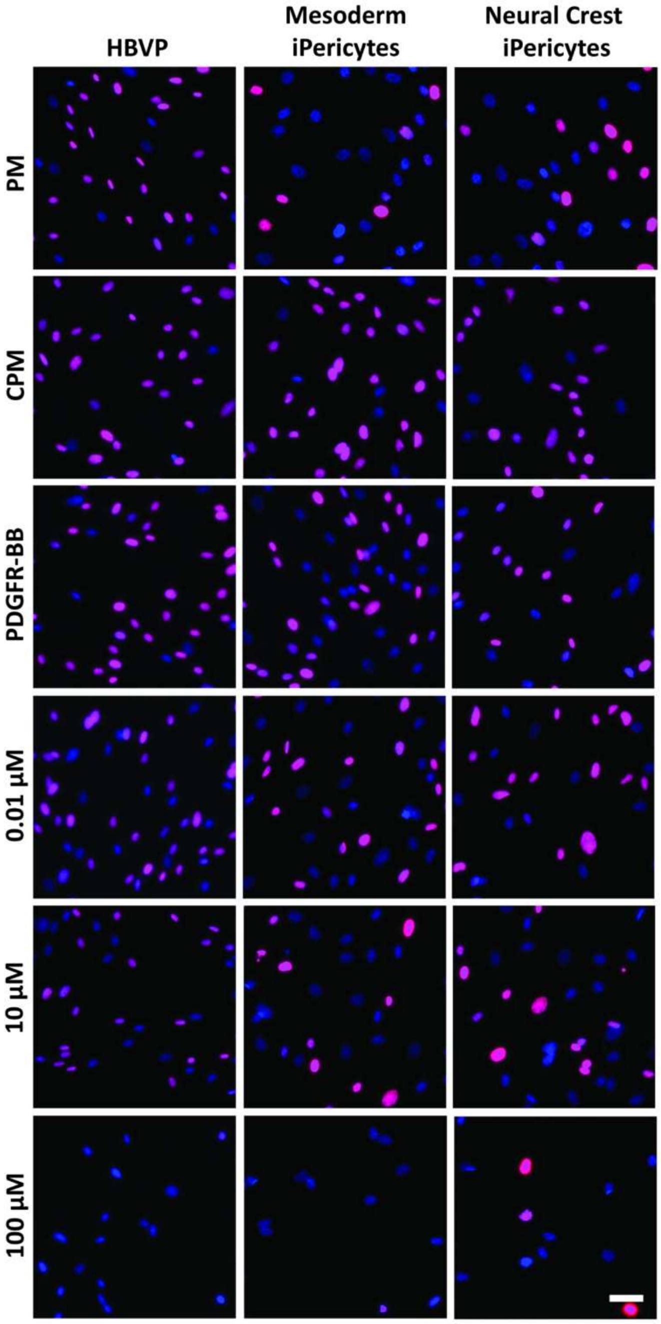


**Supplementary Figure 3. PDGF-BB and imatinib alter proliferative cell number of iPericytes *in vitro.*** Representative images of EdU (magenta) staining in HBVP, mesoderm iPericytes and neural crest iPericytes treated with basal pericyte media (PM), complete pericyte media (CPM), PM + PDGF-BB (a pericyte growth factor), and PM + PDGF-BB + increasing concentrations of imatinib (a PDGFRβ inhibitor). Total cells were identified using DAPI (blue). Scale = 50µm.

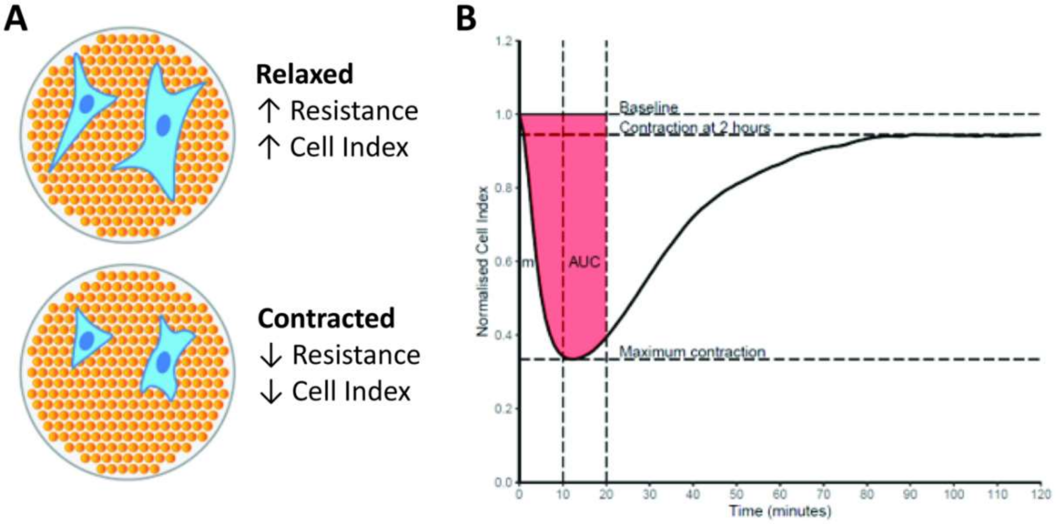


**Supplementary Figure 4. Schematic of pericyte contraction measurement and analysis using the xCelligence system.** To test contractile function of neural crest iPericytes and mesoderm iPericytes in comparison to HBVPs, the contractile response of cells was measured using an xCelligence system. (A) Cells were plated in specialised cell culture plates with electrodes which measure electron flow. Relaxed cells have greater cell volume meaning larger impedance to electron flow, thus increased resistance, which is interpreted as a higher 'Cell Index'. Contracted cells have reduced cell volume meaning less impedance to electron flow resulting in reduced resistance, which is interpreted as lower 'Cell Index'. (B) For contractile studies, cell index for each condition was measured every minute for 2h and was normalised to time point 0, when vasoactive agents were added. To quantify differences between groups, three measures were extracted from the normalised cell index over time: slope *(m)* shows the rate of contraction over the first 10 mins and is calculated as the slope of the linear equation y = mx+b for each replicate from t = 0 to t = 10; area under the curve (AUC) shows the cumulative change in contraction over time and is calculated as the area (A) under the curve between t = 0 and t = 20 mins; Δ cell index shows the change in cell index from baseline at specific time points, e.g. at the time of maximum contraction, contraction after 2h.

### Supplementary Data

The below is code used in QuPath for the analysis of the proliferation assay.

**Select all ROI. groovy**

```
createSelectAllObject(true);
```

**EDU Analysis. groovy**

```
createSelectAllObject(true);

selectAnnotations();

runPlugin('qupath.imagej.detect.cells.PositiveCellDetection', '"detectionImage": "DAPI",
"requestedPixelSizeMicrons": 0.25,  "backgroundRadiusMicrons": 8.0,
"medianRadiusMicrons": 0.0,  "sigmaMicrons": 5.0,  "minAreaMicrons": 50.0,
"maxAreaMicrons": 600.0,  "threshold": 20.0,  "watershedPostProcess": true,
"cellExpansionMicrons": 0.0,  "includeNuclei": true,  "smoothBoundaries": true,
"makeMeasurements": true,  "thresholdCompartment": "Nucleus: EDU mean",
"thresholdPositive1": 2000.0,  "thresholdPositive2": 0.0,  "thresholdPositive3": 0.0,
"singleThreshold": true');
```
